# A room-temperature ferrimagnet made of metallo-proteins?

**DOI:** 10.1101/094607

**Authors:** Michael Winklhofer, Henrik Mouritsen

**Affiliations:** Institut für Biologie und Umweltwissenschaften, Universität Oldenburg, D-26111 Oldenburg, Germany; Fakultät für Physik, Universität Duisburg Essen, D-47057 Duisburg, Germany

## Abstract

Qin et al. (A magnetic protein compass, Nature Materials 15, 217-226, 2016) claim that "MagR is the first known protein that carries an intrinsic magnetic moment at ambient temperature". We show here that the claim must, unfortunately, be fundamentally wrong.

---

Qin et al. 2015 [1] present a highly innovative strategy for finding interaction partners with the candidate magnetosensory protein cryptochrome (Cry). They rename IscA1 as MagR and present it as a putative interaction partner of Cry4, which in itself will be a truly exciting result if confirmed by other research groups. Yet, we have difficulties in comprehending at least three of the key claims in the paper: (a) That the iron-sulfur cluster protein would be so strongly magnetic at room temperature that it aligns with the Earth’s magnetic field; (b) that the interaction partner for Cry should be a strongly magnetic entity; and (c) that a magnetoreceptive Cry4 should be located in many different cell types within the retina.

To (a): The magnetization curve data (Fig. 5e and Supp. Fig. 16 of [1]] suggests a very small protein magnetization, below 10^-3^ emu/g in a magnetic field of *H=100* Oe (10 mT). A magnetization of 10^−3^ emu/g translates into a magnetic dipole moment *m_p_* of less than 1 *μ_B_* (Bohr magneton, i.e., the magnetic moment of a single spin) per Cry4-MagR complex consisting of 20 MagR and 10 Cry units with a total molecular weight of about 840 kDa. Consequently, the magnetic moment of the Isca1-Cry4 complex would according to the measurements of Qin et al.’s [1] own data on average be smaller than that of a free radical (for details, see supplement). Furthermore, for *m_p_*=1 *μ_B_* at *H*=0.5 Oe, the magnetic energy is ten million times smaller than the thermal energy at room temperature, which would imply a random distribution in an orientation experiment as shown in Fig. 5a of [1). In contrast, as shown in SI Material here, to produce the distribution of long axes shown in Fig. 5b of [1], the magnetic moment *m_p_* would have to be seven orders of magnitude higher than the value determined from the magnetization curves. Therefore, the alignment obtained in [1] must be an artifact due to some systematic bias.

We also find it hard to understand from a theoretical point of view how the claimed ferrimagnetic properties with a stable magnetic polarity at room temperature could possibly emerge in the Cry4-MagR-polymer protein complex, all the more so because electron spin resonance and Mössbauer spectroscopy have shown iron-sulfur cluster proteins to be either diamagnetic or paramagnetic [2] and that there is only one known class of organometallic magnets in which room temperature ferrimagnetism exists: the polymer ferrimagnet V^II^[TCNE]_x_ (*TCNE*=tetracyanoethylene, x≈2) and its analogs [3]. Compared to the Cry4-MagR complex, the V^II^[TCNE]_x_ and related designer magnets have abundant paramagnetic centers that are bridged in a three-dimensional network structure by cyanide ligands that mediate strong antiferromagnetic coupling between the paramagnetic centers of different spin quantum numbers and thereby cause ferrimagnetism [4]. Such a strong and three-dimensional coupling of spins is a key requirement for a high ordering temperature (i.e., room-temperature ferri/ferromagnetism). In contrast, the Cry4-MagR complex (structure shown in Fig. 3 of [1]) with its sparse iron centers obviously lacks this requirement: The few “iron loops” are too isolated in the structure, *i.e.,* too far away from adjacent iron loops to be significantly exchange coupled. The only strong exchange coupling present in the system is *within* a single [2Fe-2S] cluster, which is of antiferromagnetic nature (superexchange via sulfide bridge) and couples the 2 Fe spins antiparallel to each other, resulting either in net spin, *S*=0 (diamagnetic, if both Fe have the same valence), or a net paramagnetic spin *s*=1/2 (just like a free radical) in the mixed valence system [Fe^III^Fe^II^-2S], see [2]). Exchange interactions *among* paramagnetic [2Fe-2S] clusters of adjacent IscA1 units will be weak (i.e., dipolar-dipolar coupling instead of orbital overlap) and may lead to spontaneous magnetization below a few Kelvin, but no spontaneous magnetization at anywhere near room temperature.

To us, therefore, contamination rather than intrinsic protein magnetism seems to be the only plausible reason why the protein crystals were observed to spin in a rotating magnetic field. Although the light-microscopy based technique used does not yield false positives (because an object spinning coherently with the magnetic field must be magnetic], it requires further transmission electron microscopic analysis to rule out contamination artifacts [5].

To (b): The radical-pair mechanism does not require a magnetic protein partner to work [6], and if an interaction partner of Cry were so strongly magnetic that it aligns to the Earth's magnetic field, there would be no need for Cry as a magnetoreceptor. Of course, it is possible that not Cry4, but MagR is the magnetoreceptor and that Cry4 would only be a non-magnetically sensitive part of the signaling transduction cascade. However, that possibility is in poor agreement with several other results in the field such as the high sensitivity to radio-frequency magnetic fields observed in migratory birds [7].

To (c): The Cry4 and IscA1 stainings in Figure 4 of [1] basically report that both proteins are located in more or less all cells in the retina, and we suspect that they would be found in most cells in the rest of the nervous system, too, if the authors of [1] did the staining with their antibodies. If Cry4 and/or IscA1 were part of a magnetic sensory system, we would expect at least one of the proteins to be located only in a subset of cells forming part of a specific sensory pathway [8]. According to our experience, when an antibody stains more or less all cells in the retina, it either detects a very common housekeeping gene, which does not play a highly specific sensory role in a specific sensory system, or there is a problem with antibody specificity.

## SUPPLEMENTARY INFORMATION

Qin et al. 2016 [1] report that the putative clCry4/clMagR magnetosensor protein complex will preferentially align with the axis of an applied magnetic field when placed on an EM grid. Surprisingly, even in the comparatively weak geomagnetic field, about 45% of the isolated rod-like protein particles were found to be oriented with their long axis roughly parallel to the geomagnetic field (0.4 G or 0.04 mT), while for the null hypothesis of random orientation, all the three bins shown in Fig 5b of [1] would be expected to be equally populated with 33% each. In a 25 times stronger magnetic field, the fraction of protein particles in the “parallel bin” increased further to 55% [1]. Here, we will show why the reported 45-55% must represent an experimental artifact and show that the magnetosensor protein would need to have an extraordinarily large magnetic moment (about six orders of magnitude larger than the magnetic moment claimed to have been measured by the authors in Figure 5e and Supplementary Figure 16 of [1]) to produce the obtained orientation statistics.

The magnetization data (after subtraction of the diamagnetic background) shown in Fig. 5a of Qin et al. [1] appear too noisy to be significant. Even if we were to interpret the noise as magnetization signal due the protein, then a magnetization of at most 10^-3^ emu/gram is still at least four orders of magnitude lower than that of a proper molecular magnet with 10 - 50 emu/gram (see Table 2 of [2]). From a specific magnetization *σ_s_* =10^-3^ emu/gram (=10^-3^ Am^2^/kg in SI units), we can estimate the molar magnetization *σ_n_* of the protein complex, which consists of 20 units of Isca1 (M_Isca1_=14.5 kg/mol) and 10 units of Cry4 (M_cry4_=55 kg/mol):

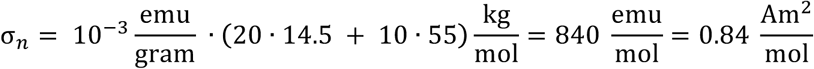

The magnetic moment per protein is obtained by division of the molar magnetic magnetization by the Avogadro number *N_A_*,

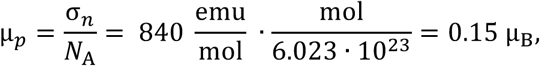

where *μ_B_* =9.274 · 10^-21^ emu is the Bohr magneton (in c.g.s. units), i.e., the magnetic moment of an electron spin. Consequently, the magnetic moment of the Isca1-Cry4 complex would on average be smaller than that of a free radical! It is clear from this elementary calculation that with such a low magnetic moment, the protein complex could not possibly align with an Earth-strength magnetic field.

In the following, we provide an order of magnitude estimate of the magnetic moment per protein needed to produce a preferential alignment as in Fig. 5b of [1]. Applying Boltzmann statistics, the probability of finding a magnetic molecule with its magnetic dipole moment vector 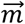 oriented at an angle *θ* relative to the magnetic field vector 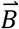 is proportional to its Boltzmann factor exp(-*E_m_*(*θ*)/*kT*), where 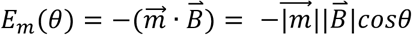 is the magnetic energy (Zeeman energy), *k =* 1.38 · 10^-23^J/K is the Boltzmann constant and *T* is the absolute temperature in Kelvin. The probabilities are normalized [3] to unity by integrating over all possible orientations in the plane, using the Boltzmann factor of each orientation as a weight function: 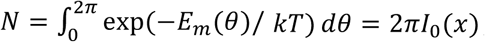, where *I*_0_ is the modified Bessel function of the first kind and 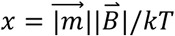 is the magnetic to thermal energy ratio.

Without loss of generality, we assume that all molecules have similar absolute value 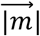 and that the vector 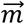 is coaxial with the long-axis of the molecule, but can have either polarity with respect to the molecule coordinate frame. Since the magnetic polarity of 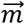 cannot be extracted from the orientation data, but only its axial orientation, we have to consider that a molecule in any given geometric orientation can actually represent either of the two magnetic polarity states (see Fig. S1 here). In the example shown, the weight for a molecule 1 in polarity state 1 (on the left in Fig. S1) with its magnetic moment 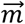 oriented at an angle *θ*_1_ < 90° is given by 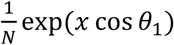. For molecule (right in Fig. S1) having polarity state 2 opposite to polarity state 1, but otherwise same axial orientation, the orientation angle of its magnetic moment is *θ*_2_ = 180° + *θ*_1_ and its weight is 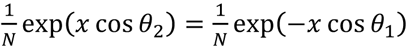. That means that the probability of finding a molecule in the favorable polarity state 1 is exp(2 *x* cos *θ*_1_) times higher than finding it in the opposite polarity state. Taken together, the two possible polarity states result in a probability density of 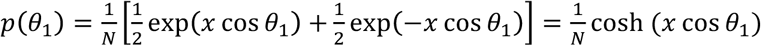 for the molecule to be oriented at an angle *θ*_1_.

**Fig. S1.**
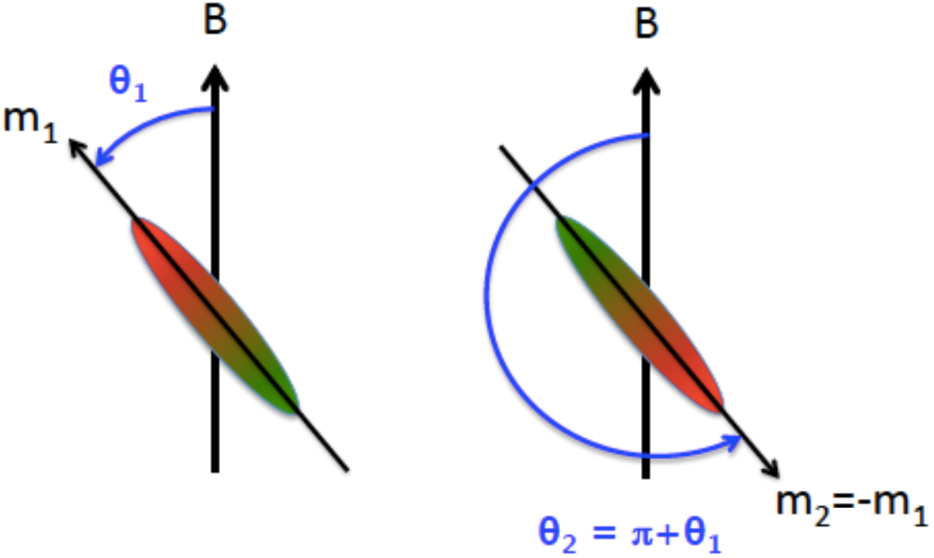
For a given axial orientation, a molecule can have either polarity state. The “up” polarity state on the left has a higher probability to occur than the “down” polarity state on the right. In case the polarity is not known, which is relevant here, both possibilities have to be considered when deriving the probability for the molecule axis to be oriented at an angle θ_1_ with respect to the external field.

Given that the magnetization curve published in Qin et al. [1] for the Isca1-Cry4 complex shows that the x-value for the Isca1-Cry4 complex is 0.0000001, there is no way to reproduce the orientation data of Fig. 5b. We can only reproduce key features of the orientation data from Fig 5b of [1] if we insert x values of about 2, i.e. ten million times larger than the value measured for the Isca1-Cry4 complex by Qin et al. [1] (see Fig. S2 here). The magnetic moment 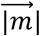 for *x*=2 with *B*=0.04 mT amounts to 0.2 fAm^2^ = 2.2 · 10^7^ *μ_B_*, where *μ_B_* is the Bohr magneton *μ_B_* = 9.274 · 10^-24^ Am^2^ in SI units). From these calculations, it is clear that the preferential axial orientation shown in [1] must be an experimental artifact. We would have liked to see control experiments conducted in a zero magnetic field, where random orientation is expected (i.e., about 33% for each of the chosen bins).

**Fig. S2.**
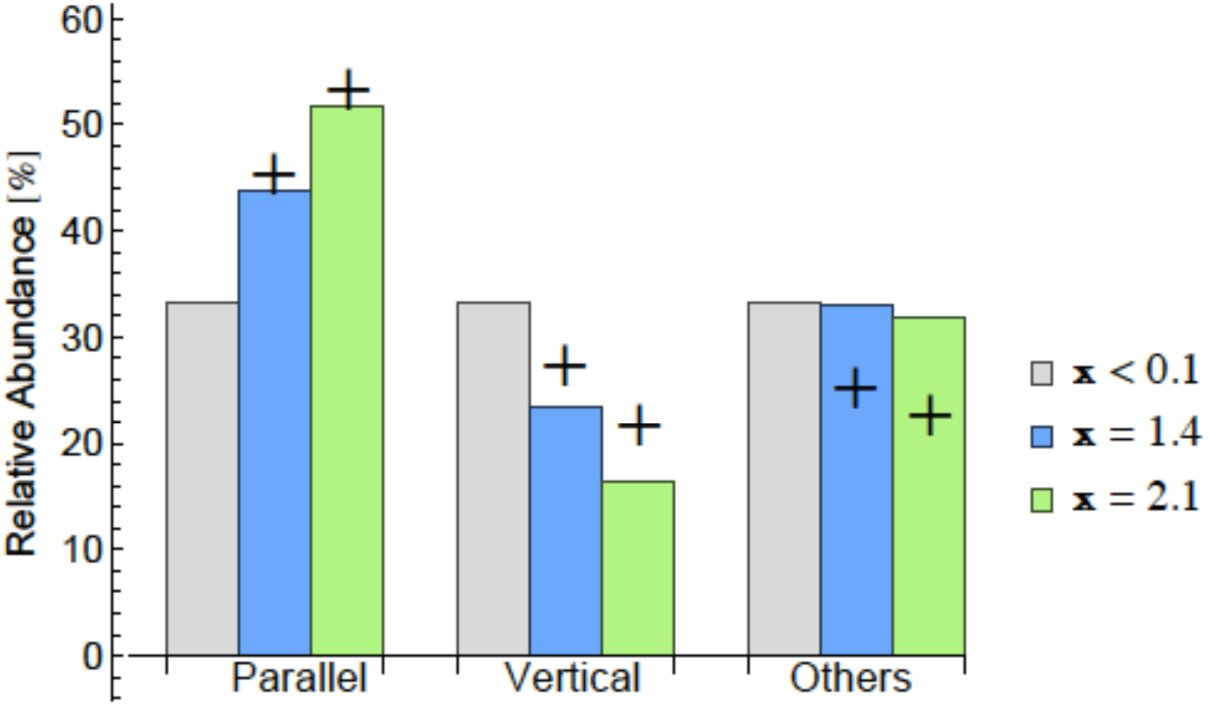
Distribution of molecule axes modeled with Boltzmann statistics using a Boltzmann factor of x<0.1 (grey), x= 1.4 (blue) for B = 0.04 mT and ofx = 2.1 (green) for B = 1 mT, in comparison with experimental data from [1] marked by + symbols.

[3] Since θ is a continuous variable, the expression 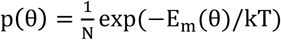 is a probability density function. The probability of vector 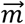 to be oriented at an angle between θ and θ + dθ, where dθ is an infinitesimal angular increment, is given by ρ(θ)dθ. While ρ(θ) can be greater than 1, ρ(θ)dθ is normalized to unity in an integral sense: 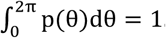 ρ(θ)dθ = 1. The probability of finding 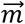 to be oriented between two macroscopic angles, say between θ = 0 and θ = π/6 is given by 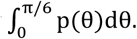

## References

[1] Qin, S., et al. Nature Materials, 14, (2015), doi: 10.1038/NMAT4484

[2] Hagen, W. R., EPR spectroscopy of iron-sulfur proteins, in Advances in Inorganic Chemistry, vol 38: Iron-sulfur proteins (ed R. Cammack], Academic Press, San Diego (1992); Cammack, R. & F. MacMillan. In Metals in Biology: Applications of High-Resolution EPR of Metalloenzymes, Biological Magnetic Resonance 29, Springer, New York (2010); Pandelia, M.E., N. Lanz, S. J. Booker & C. Krebs. Biochim. Biophys. Acta 1853, 1395–1405 (2015)

[3] Manriquez, J. M., G. T. Yee, R. S. McLean, A. J. Epstein & J. S. Miller. Science 252, 1415–1417 (1991); Thorum, M. S., K. Pokhodnya, J.S. Miller. Polyhedron 25, 1927–1930 (2006).

[4] Miller, J.S. & A.J. Epstein. Angew. Chem. Int. Ed. 33, 385–415 (1994); Verdaguer, M. & G. S. Girolami. in: Magnetism: Molecules to Materials V (eds Miller, J.S. & M. Drillon) Wiley-VCH, Weinheim (2004)

[5] Eder, S. H. K., et al., Proc. Natl. Acad. Sci. USA 109, 12022–12027 (2012); Edelmann, N. B., et al. Proc. Natl. Acad. Sci. USA 112, 262–267 (2015)

[6] Mouritsen, H. & P. J. Hore, Curr. Op. Neurobiol., 22, 343–352 (2012); Hore, P. J. & H. Mouritsen, Ann. Rev. Biophys. 45: in press (2016)

[7] Engels, S., et al., Nature 509, 353–356 (2014)

[8] Niessner, C. et al., PLoS ONE 6: e20091 (2011); Bolte et al. PLoS ONE 11: e0147819 (2016); Niessner, C. et al., Sci. Rep. 6: 21848 (2016)

## References

[1] S. Qin et al., A magnetic protein compass, Nature Materials 15, 217–226 (2016)

[2] J. S. Miller & A. J. Epstein, Organic and organometallic molecular magnetic materialsdesigner magnets, Angew. Chem. Int. Ed. 33, 385–415 (1994)

